# TissueViewer: A Web-Based Multiplexed Image Viewer

**DOI:** 10.1101/2024.06.21.600100

**Authors:** D.G.P. van IJzendoorn, M. Matusiak, R.B. West, M. van de Rijn

**Affiliations:** Stanford University, Department of Pathology; Leiden University Medical Center, Department of Pathology

## Abstract

Datasets generated by spatial biology techniques such as multiplex immunofluorescence staining or spatial transcriptomics profiling of histologic sections carry a tremendous wealth of information. Several commercial platforms exist that can simultaneously acquire 1-1000 distinct marker signals (e.g., MIBI, CODEX, Orion, Nanostring CosMX SMI, Vizgen). However, due to the large size of these datasets, their viewing and sharing are slow, laborious, and require extensive computational resources. To overcome this, we developed TissueViewer, a web-based viewer designed to deliver high-resolution images over the internet with low bandwidth requirements and at high speed.

## Main

Recent advances in spatial biology enable routine imaging of histologic specimens at sub-cellular resolution. Tens to thousands of markers can be simultaneously acquired, producing hundreds of gigabytes of data per image ^1^. Imaging instruments are bundled with proprietary software packages to allow for rapid viewing of the spatial datasets. Yet, these packages are part of the instrument used and may be accessible only on local machines associated with those instruments. In addition, due to the large size of the data, its handling often requires extensive computational resources and coding skills, and sharing datasets with colleagues in other laboratories is slow and laborious. To overcome these obstacles, we developed TissueViewer, a lightweight server-client application designed for viewing and sharing high-resolution, multi-color imaging files. The main innovation is based on a new way to deliver high-resolution spatial images stored on a server or in the cloud to a local machine using low data transfer and high speed. TissueViewer thus facilitates spatial data analysis and sharing not only within laboratories but also globally.

Software packages designed for local image viewing, such as Photoshop, QuPath, Fiji, Napari, or Mantis, require the image data to be stored locally and fully loaded into the computer’s memory. On a laptop or desktop computer, this causes the images to load slowly, freeze, and often crash the software. To alleviate this problem, researchers have to heavily downscale the data resolution and load selected channels and image regions at a time.

Sharing and viewing the content of multi-marker immunostains over the internet is currently supported by two methods. The first approach distributes imaging storage formats, such as OME-TIFF files, directly over the Internet ^2^. Applications that use this approach allow the user to select any number of channels to display; however, even though these methods retrieve only relevant fields and channels, we found that these tools still require large amounts of data to be received and read in memory by the user, resulting in long data loading times. This phenomenon arises from inherent challenges associated with retrieving precise data fragments from expansive files over an internet network, characterized by constraints such as latency and bandwidth limitations. Therefore, the OME-TIFF data sharing approach can be costly to host and slow to load for the user. One application that serves OME-TIFF directly is Aviator ^3^.

The second approach is based on Deep Zoom Images (DZI), a format developed by Microsoft ^4^, where the imaging data gets formatted into pyramidal images. This method is very efficient at delivering only those fragments of data currently being viewed by the user and therefore is very fast and requires relatively low bandwidth. The downside is that this approach requires pre-selection at the source of 3-4 channels of interest at a time without the ability to change the markers being examined. An example of a viewer using this technique is Minerva (in story mode) ^5^.

We developed an approach that builds on the favorable aspects of the two methods described above and solves their inefficiencies. Specifically, TissueViewer enables flexible viewing of any number of channels with a low amount of data transfer, low memory usage, and high speed.

TissueViewer uses a lightweight server-side function that generateds DZI images for the location and channels of interest and sends the image, in DZI format, to the client (**Fig 1A**). The result is that the user retains the flexibility to select a channel containing data for a specific region and can adjust the staining intensity level at all times. We found that this approach reduces the amount of data transfer and memory usage needed to display the image. This allows for cost-effective deployment on large cloud providers such as the Google Cloud Platform, Amazon AWS, Microsoft Azure, or the possibility of hosting Docker containers on institutional clusters. Due to the low-cost nature of this approach, it can be used to include browsable full-resolution imaging data with publications, as demonstrated in our recent study ^6^.

**Fig 1.**
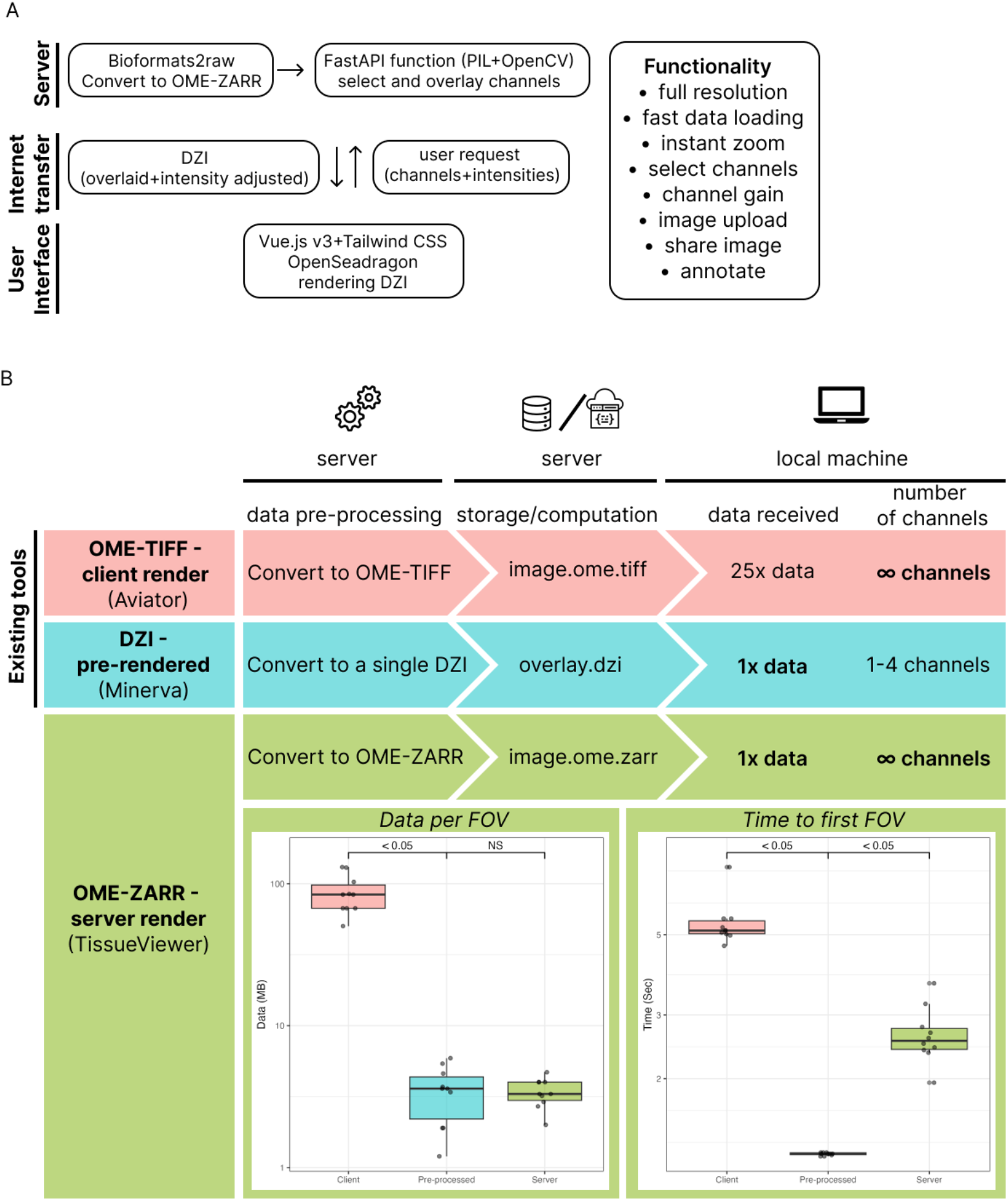
TissueViewer viewer allows for instant and intuitive browsing of spatial multiplex immunostaining. **(A)** Overview of TissueViewer, built as a client-server application. **(B)** T*op: The schematic* illustrates a comparison of two existing methods of sharing and viewing online multiplex imaging data with TissueViewer. Direct OME-TIFF file sharing (Aviator) offers significant flexibility in selecting channels containing a specific marker and adjusting staining intensity, yet is slower due to high data transfer requirements. Pyramid Deep Zoom Image (Minerva) tiling is faster and requires less data being received by the user; however, it restricts channel numbers and requires channel preselection at the source. TissueViewer pre-renders the OME-ZARR file into Deep Zoom Image tiles using a lightweight server function that requires minimal local resources. This approach combines the speed and low data usage of single Deep Zoom Images. *Bottom left:* to benchmark the performance of TissueViewer, we compared the data used to load a field of view (FOV) and *bottom right:* the time required to load the application and display the first FOV.

We compared network data usage and load times between sharing OME-TIFF, a pre-processed DZI, and our OME-TIFF/DZI approach using high-resolution multiplexed imaging data generated on the Orion Rarecyte platform as a benchmark. The files were served from the same server and hard disk, and we displayed four channels for all methods. We observed a significant reduction in bandwidth usage when using TissueViewer compared to both viewing the OME-TIFF file with Aviator viewer and as a DZI file with Minerva. Specifically, there was a 25-fold reduction in data transferred to view the image between Aviator and TissueViewer and similar data usage between Minerva and TissueViewer (**Fig 1**., *bottom left*). Load time, including loading the application and showing the first field of view, was 5.4 seconds for the OME-TIFF, 1.23 seconds for the preprocessed DZI file, and 2.65 seconds using the TissueViewer hybrid approach (**Fig 1**., *bottom right*).

TissueViewer handles both multi-channel immunofluorescence (**Fig 2**.) and single-channel histology imaging such as H&E stains. TissueViewer includes an integration with bioformats2raw ^7^ to convert files to OME-ZARR for serving, with the capability to watch folders for new files to convert, mitigating added complexity. While the server function adds some complexity, the reduced data requirements make this a minor concern since computing and storage costs are nearly identical. Moreover, the server function is lightweight and can be easily run in a serverless environment, enabling TissueViewer to potentially handle many concurrent requests. TissueViewer allows the raw files to be made available as downloadable links or upon request for further analysis. TissueViewer is easy-to-setup and allows users to extend it with additional features.

**Fig 2.**
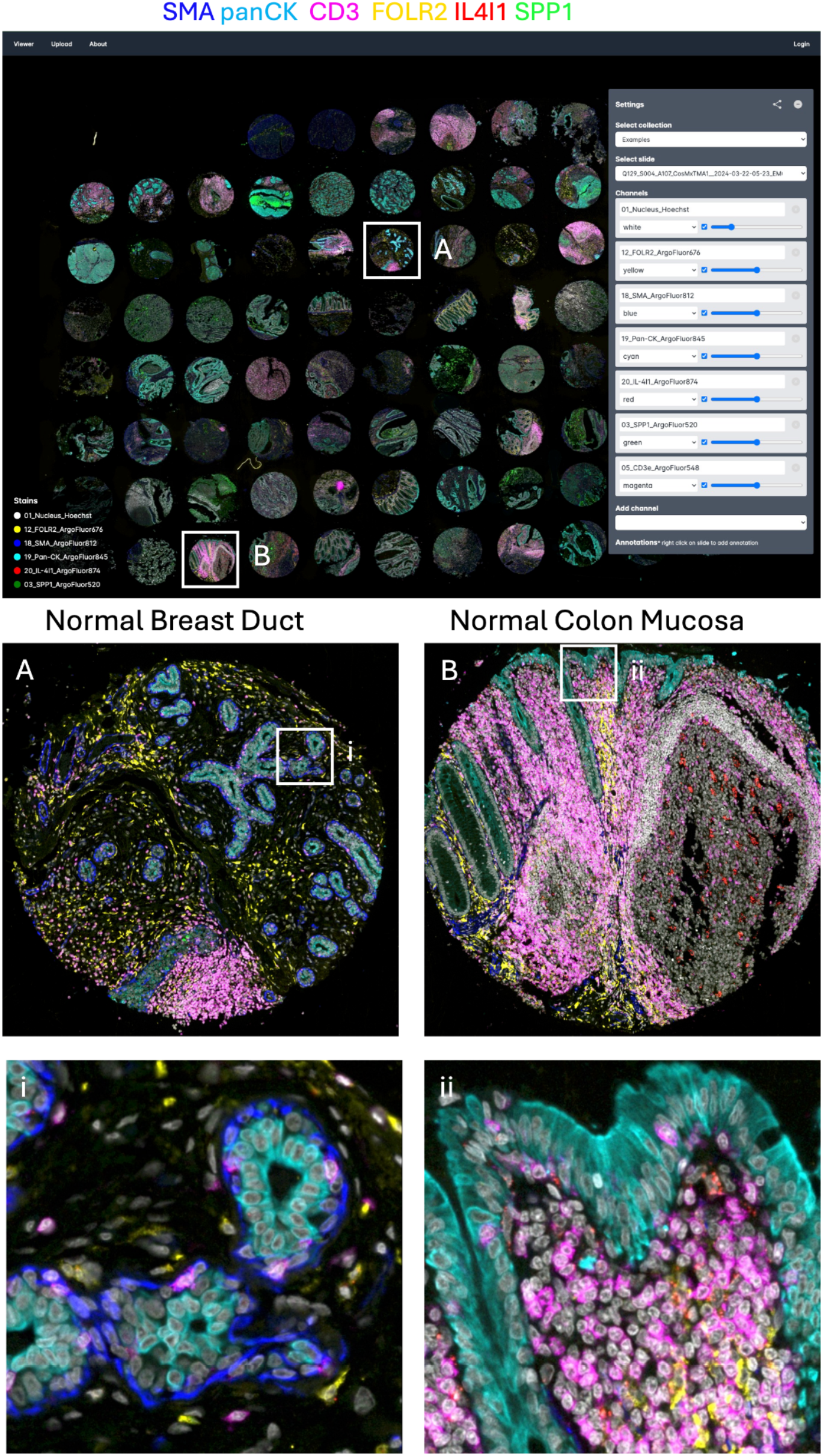
TissueViewer allows for instant high-resolution zoom. *Top:* Full tissue microarray view and TissueViewer interface, *Bottom:* Zoomed-in regions correspond to boxed regions on the top.

In conclusion, TissueViewer was developed to enable fast and easy sharing and viewing of highly multiplex imaging data. Our approach demonstrated a 25-fold lower data transfer requirement compared to sharing raw files directly. Importantly, this reduction in data transfer usage on the user side resulted in much faster image load times compared to sharing OME-TIFF data, and only a small loss in speed compared to the pre-processed DZI approach.

TissueViewer, developed as a lightweight server-client application, reduces image load times of large multiplexed images to enhance collaboration and image sharing. As the first fast and flexible method to share high-resolution multiplex imaging data, TissueViewer will democratize research by making image viewing more accessible, even with slower internet speeds.

TissueViewer is available on GitHub and can be used on the TissueViewer.org platform, where readers can upload their own data with a limit of 50 GB to share with colleagues.

## Methods

### Application design

TissueViewer is a server-client application that includes an automated pipeline for data preparation. This pipeline directs files to an ingestion folder, where newly added multiplex imaging files are automatically processed. Utilizing the bioformats2raw, multi-channel imaging data is converted to OME-ZARR files.

On the server side, a FastAPI-based Python application manages API calls from the client, which specifies the desired channel colors and intensities. Upon receiving a request, the server retrieves the corresponding image channels using the Dask Array python package. It then mixes the channels using OpenCV, substantially reducing the data volume transmitted to the client.

The client interface is developed in Vue.js v3, employing Tailwind CSS for UI components and OpenSeaDragon to render the DZI images.

Designed for seamless cloud deployment, DZI images can be stored in services like Google Cloud Buckets, and the server can be hosted on Google Cloud Run. Our GitHub repository provides detailed deployment guidelines.

### Benchmark TissueViewer

To compare performance and data usage between TissueViewer and other tools that directly access OME-TIFF files (such as Aviator), we used an updated Chrome browser and the Network analysis developer console to measure data usage in megabytes (MB) and load times in seconds. The same imaging file was opened in TissueViewer (processed as DZI channels) and as a pre-processed DZI file in OpenSeaDragon, and we compared the data usage when opening the OME-TIFF file using Aviator. The images were hosted on the same hard drive in an Ubuntu 20.04 Server over the internet. All viewers were set to load the same 10 areas at the same zoom levels.

### Expandability

TissueViewer was built using open-source components, making it easily expandable with additional features. This also opens up the possibility to add analysis pipelines and additional collaboration features.

### Immunofluorescence imaging

The slides were stained with a cocktail of antibodies conjugated to ArgoFluor dyes according to the manufacturer’s guidelines and scanned on the Orion v.2 instrument.

## Acknowledgements

This research was supported by an unrestricted grant of Stichting Hanarth Fonds, The Netherlands to DVIJ. This research was supported by the Virginia and D.K. Ludwig Fund for Cancer Research and the Taube Family Foundation.

